# A comparison of reference-based algorithms for correcting cell-type heterogeneity in Epigenome-Wide Association Studies

**DOI:** 10.1101/101709

**Authors:** Andrew E Teschendorff, Charles E Breeze, Shijie C Zheng, Stephan Beck

**Affiliations:** CAS Key Lab of Computational Biology, CAS-MPG Partner Institute for Computational Biology, Shanghai Institute for Biological Sciences, Chinese Academy of Sciences, Shanghai 20031, China.; Department of Women‘s Cancer, University College London, 74 Huntley Street, London WC1E 6AU, United Kingdom.; Statistical Cancer Genomics, Paul O’Gorman Building, UCL Cancer Institute, University College London, 72 Huntley Street, London WC1E 6BT, United Kingdom.; Medical Genomics, Paul O’Gorman Building, UCL Cancer Institute, University College London, 72 Huntley Street, London WC1E 6BT, United Kingdom.; University of Chinese Academy of Sciences, 19A Yuquan Road, Beijing 100049, China.

**Keywords:** Cellular Heterogeneity, DNA methylation, EWAS

## Abstract

**Background:** Intra-sample cellular heterogeneity presents numerous challenges to the identification of biomarkers in large Epigenome-Wide Association Studies (EWAS). While a number of reference-based deconvolution algorithms have emerged, their potential remains underexplored and a comparative evaluation of these algorithms beyond tissues such as blood is still lacking.

**Results:** Here we present a novel framework for reference-based inference, which leverages cell-type specific DNAse Hypersensitive Site (DHS) information from the NIH Epigenomics Roadmap to construct an improved reference DNA methylation database. We show that this leads to a marginal but statistically significant improvement of cell-count estimates in whole blood as well as in mixtures involving epithelial cell-types. Using this framework we compare a widely used state-of-the-art reference-based algorithm (called constrained projection) to two non-constrained approaches including CIBERSORT and a method based on robust partial correlations. We conclude that the widely-used constrained projection technique may not always be optimal. Instead, we find that the method based on robust partial correlations is generally more robust across a range of different tissue types and for realistic noise levels. We call the combined algorithm which uses DHS data and robust partial correlations for inference, EpiDISH (Epigenetic Dissection of Intra-Sample Heterogeneity). Finally, we demonstrate the added value of EpiDISH in an EWAS of smoking.

**Conclusions:** Estimating cell-type fractions and subsequent inference in EWAS may benefit from the use of non-constrained reference-based cell-type deconvolution methods.

## Background

One of the great challenges facing the omics community is that posed by intra-sample heterogeneity (ISH) [1]. Molecular profiles derived from complex tissues such as blood or breast represent averages over many different cell types. Since cell-type composition of complex tissues varies in response to phenotypes such as cancer or age, correcting for changes in tissue composition could be crucial if one wishes to identify potentially causal alterations in individual cell-types [2, 3]. This is particularly relevant for Epigenome-Wide Association Studies (EWAS) where the effect sizes of interest could be small compared to changes in tissue composition [3, 4]. Because of this, a number of different ISH deconvolution algorithms for genome-wide DNA methylation data have recently been proposed [5-9]. As with the analogous tools developed for mRNA expression data [10, 11], these algorithms can be classified as either “reference-free” [6, 8] or “reference-based” [5], depending on whether they use an a-priori database of cell-type specific DNA methylation (DNAm) reference profiles to perform the deconvolution. Although reference-free methods have the advantage that they don’t require such a DNAm database and are therefore applicable, in principle, to any tissue-type, these algorithms have only been tested in blood and don’t provide sample-specific absolute estimates of cellular proportions. Obtaining absolute estimates of underlying cell-type proportions in a tissue is an important task, as shifts in specific cell subtypes within the tissue could be used for diagnostic or prognostic purposes [12, 13], as well as providing useful mechanistic insight into systems medicine [14]. A further problem with reference-free methods is that they often rely on the assumption that the top components of variation correlate with cell-type composition, an assumption which may not always hold, and which if violated could lead to loss of biological signal. Indeed, a clear example of this can be seen in a study which used a reference-free method called EWASher [6], concluding that only a handful of CpGs are differentially methylated between breast cancer and normal breast tissue, clearly contradicting all the available evidence that DNA methylation differences between breast cancer and normal tissue is widespread and largely independent of changes in tissue composition [15, 16]. Thus, reference-based approaches appear to be the safest option provided one can construct a reference DNAm database for the tissue of interest. As representative DNAm profiles for all human normal cell types accrue [17], reference-based methods are therefore more likely to become the framework of choice for tackling the ISH problem.

So far however, only one reference-based algorithm (the Houseman algorithm) has been proposed for DNAm data [5] (see also [18, 19]). Houseman’s algorithm performs inference using a quadratic programming technique known as linear constrained projection (CP), where non-negativity and normalization constraints on cellular proportions are imposed during inference. Interestingly, in the gene expression context, a recent comparative study concluded that CP is outperformed by a non-constrained reference-based approach which relies on the technique of Support Vector Regressions (SVR) [10]. Another technique based on robust partial correlations (RPC) [10] has also not yet been explored in the context of DNA methylation data. Thus, there is an urgent need to conduct a comprehensive comparative evaluation of different reference-based deconvolution algorithms. Here, we perform such a comparison using experimental as well as computationally generated cellular mixtures, and including epithelial cell-types as well as leukocytes.

The importance of the reference database itself for the quality of the inference has also been noted before [10, 20]. Motivated by this, we here present a novel approach for the construction of a reference DNAm database, which integrates prior biological knowledge of cell-type specific sites with cell-type specific DNA methylation. Specifically, given that DNAse Hypersensitive Sites (DHS) are highly cell-type specific [21], we use such DHS data from the NIH Epigenomics Roadmap [17] and ENCODE [22, 23], to improve the quality of the reference database. Sample-specific cell-type proportions can then be estimated using either Houseman’s CP algorithm or a non-constrained approach such as SVR or RPC.

We show that incorporation of such prior biological knowledge can improve inference, and that Houseman’s algorithm is only optimal in the scenario of very noisy data. For realistic noise levels, we discover that non-constrained methods like RPC generally perform better. Hence, we propose an improved novel framework for reference-based inference of cell-type composition, called EpiDISH (<u>Epi</u>genetic <u>D</u>issection of <u>I</u>ntra-<u>S</u>ample-<u>H</u>eterogeneity), which uses (i) DHS data from the NIH Roadmap and ENCODE to construct a reference DNAm database, and (ii) RPC for estimation of cell-type proportions.

## Methods

### Reference-based algorithms for deconvolution of intra-sample heterogeneity

We considered a total of 4 different reference-based algorithms. Reference-based means that each of these algorithms models a DNAm profile of any given sample as a linear combination of a given set of reference DNAm profiles representing underlying cell-types present in the sample. Given a number *C* of underlying cell-types, each with a DNAm profile ***b***_*c*_, and denoting by ***y*
** the DNAm profile of a given sample, the underlying model is

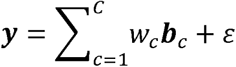

The general idea is to estimate the weight coefficients in a least squares sense, but restricting to a subset of CpGs that are highly discriminative of the underlying cell subtypes. Assuming that the reference database contains the major cell-types present in the sample ***y***, one may assume that

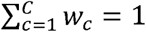

 (or more generally that

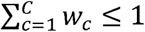
). The 4 algorithms differ in how the normalization constraint is implemented:

For 3 algorithms, the constraint that weights have to be non-negative and add to 1, is implemented *a posteriori*, i.e after inference of the weights themselves. Specifically, we follow the procedure implemented in [10] and set any negative weight estimates to zero, followed by a scaling to ensure that the non-zero positive weights add to 1. The 3 algorithms which enforce the constraints *a posteriori* are (i) multivariate linear regression or partial correlations (LR), (ii) robust multivariate linear regression or robust partial correlations (RLR/RPC) and (iii) Support Vector Regressions (SVR), an advanced form of robust penalized multivariate regression. In the case of SVR, we used the implementation called CIBERSORT [10]. For LR and RLR/RPC we used the *lm* and *rlm* R-functions (www.r-project.org), to perform the multivariate regressions.

The 4^th^ algorithm performs the inference of the weights in a least squares sense but imposes the positivity and normalization constraints as part of the inference process. This technique is known as linear constrained projection (CP) and weights can be inferred using quadratic programming (QP) [18, 19]. In implementing CP/QP there are in principle two options for the normalization constraint: one can implement a strict equality which requires the weights to add to 1, or one can implement the normalization as an inequality constraint, in which case the weights are only required to add to a number less or equal to 1. Here we implement the CP algorithm using the normalization as an inequality constraint. In effect, modulo the reference database, this algorithm is the reference-based Houseman algorithm [5]. Differences between the two implementations of CP are relatively minor since in this work we evaluate methods in tissues where all the major cell subtypes are known and for which reference DNAm profiles exist.

### Construction of integrated DHS reference DNA methylation databases

Below we give a brief summary of the datasets used in the construction of the reference databases (see also **Table 1**).

**Table 1:** Main Illumina 450k DNAm datasets used. We list the main datasets used in this study, the cell-types/tissue profiled, whether the data was used for reference database construction (if yes, we specify which cell-types were used), whether the data was used for validation/evaluation purposes (if yes, we specify which cell-types were used) and the reference/citation. Abbreviations: DNAm=DNA methylation, WB=whole blood, PBMC=peripheral blood mononuclear cells, HMEC=human mammary epithelial cells, HRCE=human renal cortical epithelia, IMR90 (fetal lung fibroblast), SCM2=Stem-Cell-Matrix Compendium-2, DMCs=differentially methylated CpGs, NK=natural killer cells, B=B-cell, Monoc=Monocytes, Neutro.=Neutrophils, Eosino=Eosinophils, CD4T=CD4+ T-cells, CD8T=CD8+ T-cells.

| Dataset Name | Tissue/cell-types | Use in Reference DNAm Database | Testing/Evaluation | Reference |
| --- | --- | --- | --- | --- |
| Reinius et al | WB, PBMC, NK, B, CD4T, CD8T, Monoc, Neutro, Eosino. (n=6 of each) | NK, B, CD4T, CD8T, Monoc., Neutro., Eosino. | WB & PBMC | [24] |
| Liu et al | WB (n=335 controls, n=354 rheumathoid arthritis cases) | No | Average Flow Cytometry estimates for cases and controls | [2] |
| Koestler et al | WB (n=18) | No | 12 Reconstructed WB mixtures + 6 WB samples with Flow Cytometry estimates | [20] |
| Zilbauer et al | PBMC, CD4T, CD8T, NK, B, Monoc, Neutro. (n=6 of each) | No | In-silico mixtures of purified blood cell subtypes | [27] |
| ENCODE | Various | HMEC, HRCE, IMR90, Liver | No | [22] |
| Slieker et al | Various | Pancreas | Liver | [26] |
| SCM2 | Various | No | HRCE, Pancreas, IMR90 | [29] |
| Lowe et al | Various | No | HMEC | [28, 35] |
| Teschendorff et al | WB (n=152) | No | Smoking associated DMCs | [31] |

#### Blood tissue

In the case of blood tissue we used the purified blood cell Illumina 450k data from Reinius et al [24]. Specifically, we used the purified cell data of Monocytes, Neutrophils, Eosinophils, CD4+ T-cells, CD8+ T-cells, Natural Killer (NK) cells and B-cells. There were 6 samples for each cell-type coming from 6 different individuals. We used a well-known empirical Bayes framework of moderated t-statistics [25] to derive differentially methylated CpGs (DMCs) between one of the 7 cell types and the rest using a false discovery rate (FDR) threshold of 0.05. Separately to this, we also identified all Illumina 450k probes that mapped to a DNase Hypersensitive Site (DHS) in any of the considered blood cell subtypes using data from the NIH Epigenomics Road map. DHS data was available for Monocytes, B-cells, T-cells and NK-cells. For each cell-type we then filtered DMCs to include only those mapping to a DHS, which we call DHS-DMCs. This resulted in 14105 B-cell, 7723 NK-cell, 12118 CD4+ T-cell, 38131 CD8+ T-cell, 11289 Monocyte, 2375 Neutrophil and 11515 Eosinophil DHS-DMCs. We then ranked these DHS-DMCs according to the mean difference in DNAm, thus favouring DHS-DMCs with large mean differences (i.e. delta beta-value > 0.8). For each cell-type we picked the top 50 DHS-DMCs. Across all 7 cell subtypes, there were 333 unique DHS-DMCs. DNAm reference centroids were then calculated as the average over the 6 samples of each purified blood cell subtype and for each of these 333 CpGs, resulting in a blood DNA methylation reference database of 333 DHS-DMCs and 7 blood cell subtypes.

#### Mixed epithelial cells

As a means of testing the statistical algorithms in an epithelial context, we sought to identify at least 3 human epithelial cell subtypes for which Illumina 450k DNAm data was available, generated as part of at least 2 independent studies, in order to use one study for reference database construction and another for validation (generation of in-silico mixtures). In addition, we also required DHS data from ENCODE or the NIH Epigenomics Roadmap for these cell-types. We identified DHS data from the NIH Roadmap for breast and pancreatic cells, whilst from ENCODE we obtained DHS data for human renal cortical epithelial (HRCE) cells. For human mammary epithelial cells (HMECs) and HRCE cells, Illumina 450k data was available from ENCODE, whereas for pancreas we used corresponding 450k data from Slieker et al [26]. In this case, because in some cases we only had 1 representative sample for each cell-type, we selected DMCs according to a difference in DNAm beta-value being larger than 0.9. This differential DNAm analysis was only performed for CpGs that mapped to a DHS in either HMECs, HRCEs or pancreas. Thus, this procedure is extremely stringent as we select DHS CpGs whose DNAm varies as much as possible between cell-types. This resulted in 272 DMCs between HMECs and HRCEs, 315 CpGs between HMECs and pancreas, and 54 DMCs between HRCEs and pancreas. In total, there were 575 unique DMCs, resulting in a DNAm reference database of 575 CpGs and 3 cell-types (breast, HRCEs and pancreas).

#### Mixed epithelial and non-epithelial cells

We identified DHS data from the NIH Roadmap for a fetal lung fibroblast cell-line (IMR90) and for B-cells, whilst from ENCODE we obtained DHS data for hepatocytes. For IMR90 and hepatocytes Illumina 450k data was available from ENCODE, whereas for B-cells we used the data from Reinius et al [24]. In this case too, because in some cases we only had 1 representative sample for each cell-type, we selected DMCs according to a difference in DNAm beta-value being larger than 0.9. This differential DNAm analysis was only performed for CpGs that mapped to a DHS in either hepatocytes, IMR90 or B-cells. Thus, this procedure is extremely stringent as we select DHS CpGs whose DNAm varies as much as possible between cell-types. This resulted in 181 DMCs between IMR90 and liver, 405 CpGs between IMR90 and B-cells, and 482 DMCs between liver and B-cells. In total, there were 912 unique DMCs, resulting in a DNAm reference database of 912 CpGs and 3 cell-types (IMR90, liver and B-cells).

### Non-DHS based reference databases

For the two previously described scenarios, we also derived reference DNAm databases without restriction to DHSs, but ensuring that reference databases were of approximately the same size as the DHS-based ones. In the case of breast, pancreas and HRCEs, this resulted in a 558 DMC x 3 cell-type reference DNAm matrix. In the case of IMR90, liver and B-cells, the reference DNAm matrix was of dimension 975 DMCs and 3 cell-types. In the case of blood, the corresponding “non-DHS” reference database was defined over 339 DMCs.

### Validation datasets

In what follows we briefly describe the Illumina 450k datasets used to validate and compare the different algorithms (see also **Table 1**).

#### Blood tissue

One dataset profiling whole blood for over 650 samples (encompassing both rheumathoid arthritis (RA) cases and controls) was available from Liu et al [2]. For this dataset, there were average flow-cytometric estimates for all major blood cell subtypes for RA cases and controls. Another dataset (Koestler et al) profiled 6 whole blood (WB) and 12 experimentally reconstructed “whole blood” mixtures [20]. In the former case, flow-cytometric estimates for the different blood cell subtypes were available for each of the 6 WB samples. In the latter case, the mixing proportions were determined by the experimentalist and therefore known without error. We also considered an additional Illumina 450k dataset from Zilbauer et al, which profiled 5 blood cell subtypes (Monocytes, Neutrophils, B-cells, CD4+ and CD8+ T-cells) with 6 replicates of each [27].

#### Other cell types

For validating and assessing the algorithms in the context of the mixed epithelial cell type scenario, we used as validation, Illumina 450k DNAm data for HMECs from Lowe et al [28], and Illumina 450k data for HRCEs and adult male & female pancreas from the Stem-Cell-Matrix Compendium-2 (SMC2) [29]. For the case of the mixed epithelial/non-epithelial cell types, we used Illumina 450k DNAm data of IMR90 (fetal lung fibroblast) from SCM2, for liver cells from Slieker et al [26], and purified B-cells from Zilbauer et al [27].

### Validation and evaluation strategy based on in-silico mixtures

For evaluation and comparison of the statistical algorithms, we generated 100 different in-silico mixtures of the purified cell DNAm profiles, with weights chosen randomly from a uniform (0,1) distribution, subject to the constraints that weights add to 1. Performance of each algorithm was then assessed using the root mean square error (RMSE) between the estimated and true weights for each cell-type, as estimated over the 100 different mixtures. R^2^ values between estimated and true weights for each cell-type were also computed over the 100 different mixtures. Average RMSE and R^2^ values over cell-types were also calculated. Finally, this procedure was repeated for a total of 25 Monte Carlo runs, yielding a total of 25 average RMSE and R^2^ values. This overall strategy was used for validation/testing purposes in the Zilbauer dataset, as well as for testing the methods in the mixed epithelial and mixed epithelial/non-epithelial scenarios.

### Noise Analysis

We note that in this work we always test methods on data which is independent from the data used to construct the reference database. This already implicitly assesses the performance of the algorithms under noise levels which one may encounter between two similar experiments performed by different labs and personnel. However, in order to assess the algorithms under increasing levels of noise, we also added Gaussian Noise of increasing variation to the mixtures. The addition of Gaussian Noise was done in the M-value basis, with data subsequently transformed back to the beta-valued basis for application of the algorithms. Specifically, we considered 7 increasing levels of standard deviation noise: SD=0, 1, 2, 3, 4, 5, For a DMC that differs between 2 cell-types by an amount of 0.8 (in a beta-valued basis), this corresponds roughly to a difference in the M-value basis of approx. 6. Thus, the case SD=6, can be seen as an extreme case of noise. The case SD=1 corresponds to a typical deviation in the beta-value basis of approximately β(1-β)/(1+β), i.e. an approx. 5% change for a CpG which is say unmethylated (β=0.05), and an approx. 16% change for a CpG which is partially methylated (β=0.4). Thus the case SD=1 is a fairly realistic scenario given known noise levels in Illumina 450k data.

### Definition of a gold-standard list of smoking-associated DMCs and of a true negative list

Smoking and whole blood is the ideal scenario in which to compare methods for cell-type correction, since many whole blood EWAS have reported strong consistency of smoking-associated DMCs (reviewed in [30]). Hence, to define a gold-standard list of sDMCs and associated genes, we used the curated table of Gao et al [30]. Specifically, we defined a gold-standard list of sDMCs defined by Illumina 450k probes which have been associated with smoking in at least 3 independent studies. This resulted in a total of 62 gold-standard sDMCs, implicating a total of 15 unique genes. Using this gold-standard list to declare a list of true positives, we then assessed sensitivity of the methods in an independent whole blood EWAS of 152 samples [31]. We note that this EWAS was not among the ones reviewed by Gao et al and hence is truly independent.

Definition of a true negative CpG (i.e. one not associated with smoking) is much harder. However, to construct an approximate set of true negative CpGs, we used the intersection of 450k probes not associated with smoking in 3 independent EWAS studies. These EWAS studies were a (i) whole blood set of 464 samples from Tsaprouni et al [32] (GEO: GSE50660), (ii) a set of 333 whole blood samples from healthy controls and (iii) corresponding 354 whole blood sample from rheumathoid arthritis cases, all from Liu et al (GEO: GSE42861) [2]. For all 3 sets, smoking status information was available. In the case of Tsaprouni et al, processed data was downloaded from GEO. In the case of Liu et al, we processed raw data with minfi[33]. All 450k data was corrected for type-2 probe bias using BMIQ [34]. P-values of association with smoking (treated as ordinal variable, 0=non-smoker, 1=ex-smoker, 2=current-smoker) was determined by linear regression in each set. We then defined CpGs as not associated with smoking if their P-value > 0.25 in each of the 3 datasets, resulting in 89290 true negative (TN) CpGs.

### Software Availability

The blood reference DNAm database for 333 DHS-DMCs and 7 blood cell subtypes is provided (**table S1 in Additional File 1**). A user-friendly R-script implementing EpiDISH is available (**Additional File 2**). EpiDISH is also freely available as an R-package from *github: https://github.com/sjczheng/EpiDISH*

## Results

### The EpiDISH algorithm and validation in blood using flow-cytometry

We collected DHS and Illumina 450k DNA methylation data from the NIH Epigenomics Roadmap [17] and ENCODE [22, 23], as well as from other Illumina 450k studies profiling individual cell-types [24, 29] and normal tissues [26, 35] (**Table 1**). Given a tissue of interest, and with prior knowledge of which cell subtypes might be present in the tissue, EpiDISH first constructs a DNA methylation reference database, by integrating DHSs from the corresponding cell subtypes with a supervised selection procedure, to identify cell subtype specific differentially methylated CpGs (DMCs) which localize to open chromatin (DHS-DMCs, **Methods, Fig.1A**). Once the reference database is constructed, EpiDISH then infers sample-specific cellular proportions using robust partial correlations (**Fig.1B, Additional File 2**).

**Figure-1.**
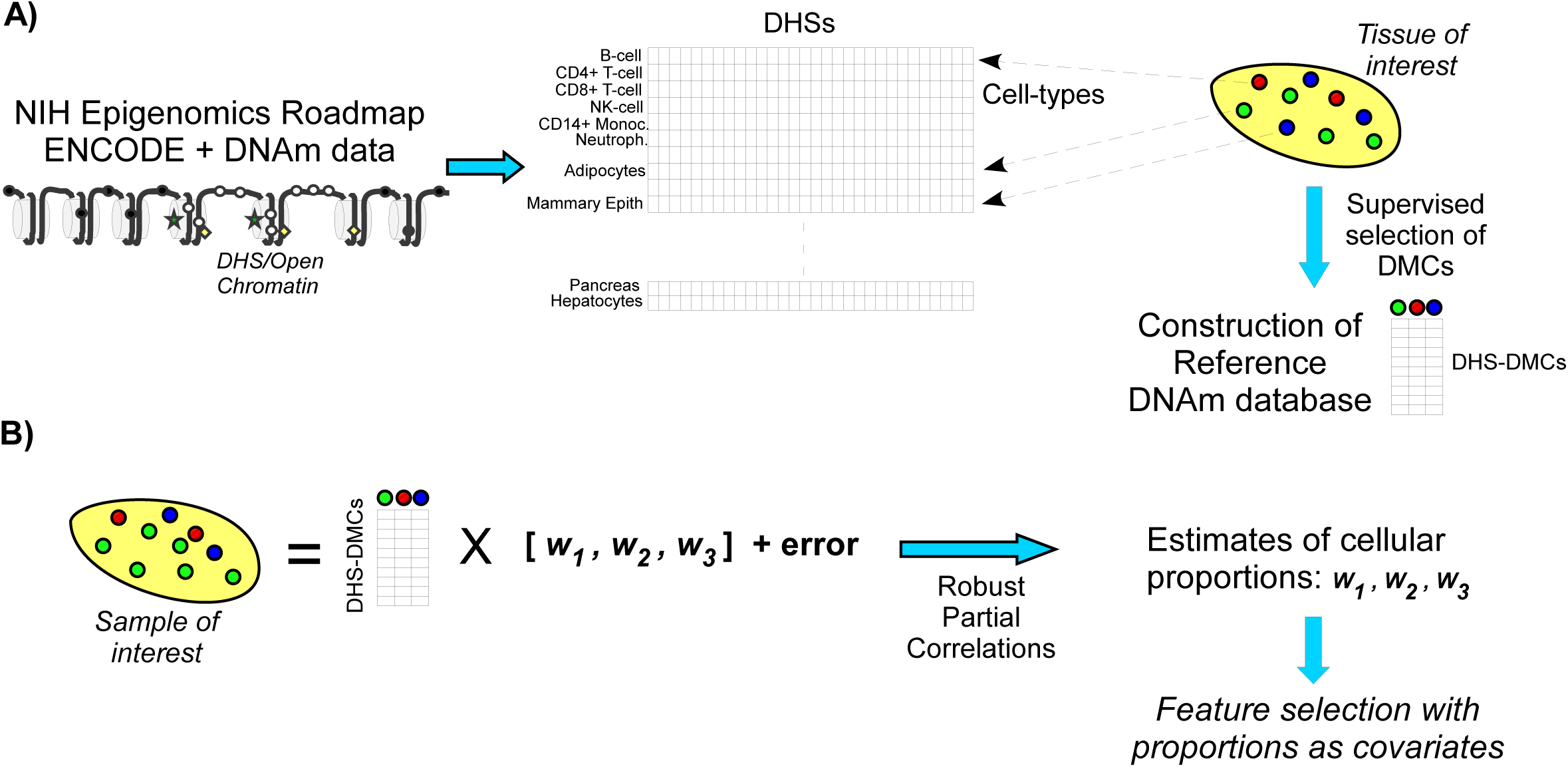
Epigenetic Dissection of Intra-Sample Heterogeneity: the EpiDISH algorithm. **A)** Given a tissue of interest and with knowledge of the main underlying cell subtypes, EpiDISH constructs a reference DNA methylation (DNAm) database for these cell subtypes using (i) DNase Hypersensitive Sites (DHS) and DNAm data for these cell types from existing public databases and (ii) a supervised selection procedure which identifies differentially methylated CpGs (DMCs) among each pair of cell-types. The resulting reference DNAm database is defined only over DMCs that map to a DHS in at least one of the underlying cell-types. **B)** Given a tissue sample of interest, EpiDISH next infers underlying cell type proportions/weights using robust partial correlations, with non-negativity and normalization constraints imposed a-posteriori. Having estimated cellular proportions for each sample, feature selection against a phenotype of interest is then performed using these proportions as covariates.

We first considered the case of blood tissue, a complex tissue for which the main constituent cell types are well known and which is being used extensively in EWAS [2]. We constructed a blood reference database using Illumina 450k DNAm profiles for a total of 7 purified blood cell subtypes (B-cells, NK-cells, CD4+ T-cells, CD8+ T-cells, Monocytes, Neutrophils and Eosinophils) obtained from [24], integrating it with blood cell subtype specific DNAse Hypersensitive Sites (DHS), as obtained from the NIH Epigenomics Roadmap [36], and further filtering the 450k probes for differential methylation between every pair of blood cell subtypes, resulting in an integrated blood reference DNAm database of 333 CpGs and 7 blood cell subtypes (**Methods**, **Table 1**, **table S1 in Additional File 1**). As a sanity check, we verified that the original purified samples segregated according to cell subtype when clustered over these 333 CpGs (**fig.S1 in Additional File 1**). Blood cell subtype proportions obtained with EpiDISH on whole blood (WB) and peripheral blood mononuclear cells (PBMC) from the same study were also in line with known proportions, i.e. strongest enrichment for neutrophils in WB and with lymphocytes making the dominant component of PBMCs (**fig.S2 in Additional File 1**).

In order to validate the EpiDISH algorithm, we first applied it to an independent 450k data set of 689 whole blood samples from an EWAS in Rheumatoid Arthritis, for which FACS (flow-cytometric) estimates of 5 purified blood cell subtypes (B-cells, NK-cells, T-cells, Granulocytes and Monocytes), averaged separately over 354 cases and 335 controls, was available [2]. We observed excellent agreement between EpiDISH and FACS estimates, for both cases and controls, with a root mean square error (RMSE) of only 4% (**Fig.2A**). Agreement was even better for the actual difference in mean blood cell subtype proportions between cases and controls, with a RMSE of 1% (**Fig.2A**). To further validate EpiDISH, we applied it to another set of 6 whole blood samples, for which independent flow-cytometry estimates of blood cell subtype fractions was available [20]. In this set too, EpiDISH achieved a RMSE of approximately 3 to 4%, with a reasonably high average R^2^ value of 0.85 (**Fig.2B**).

**Figure-2.**
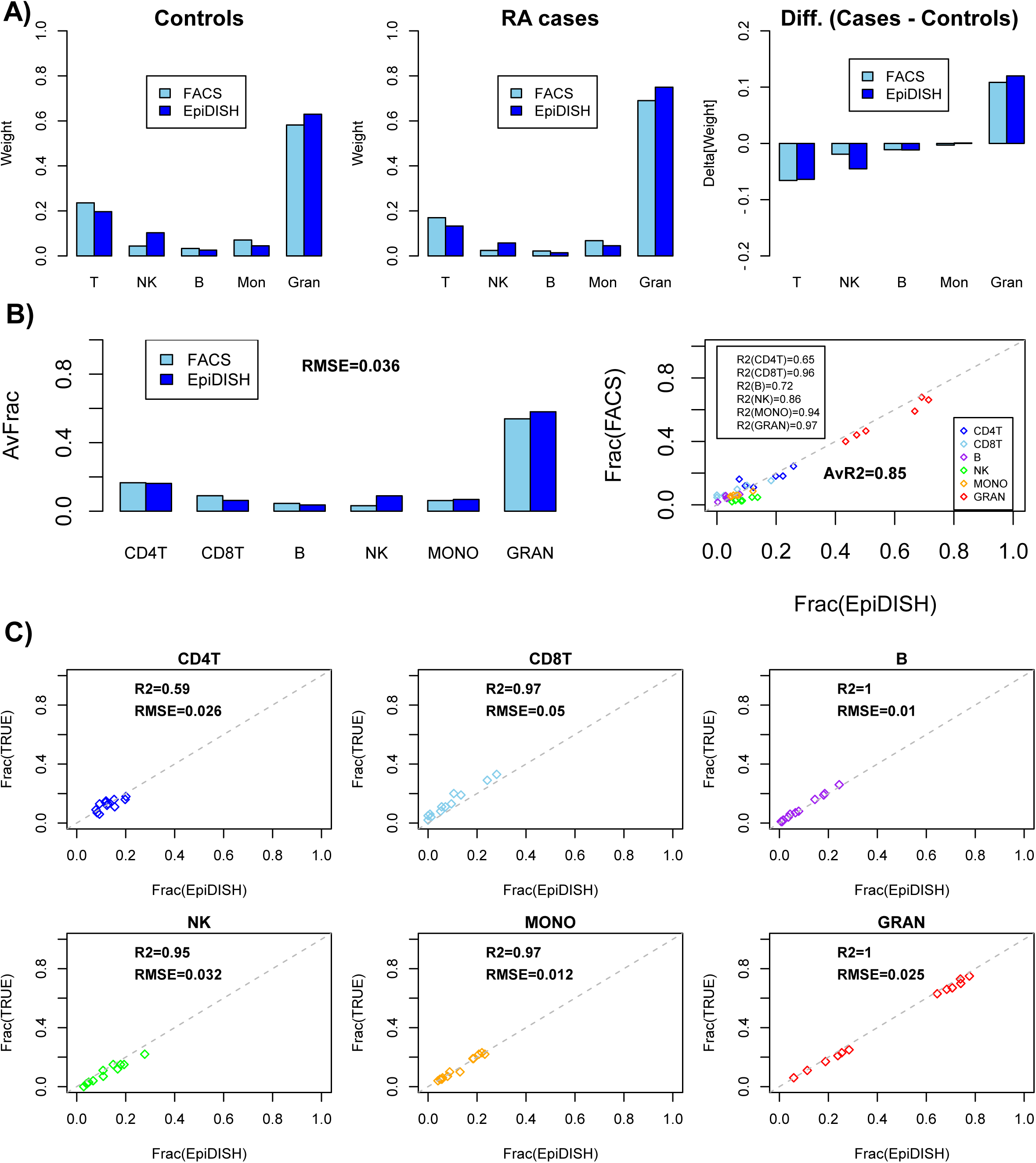
Validation of EpiDISH in independent data with FACS and known mixture estimates. **A)** Left panels: Barplots of the average weight proportions of each major bloodcell subtype according to FACS and EpiDISH, in 354 rheumatoid arthritis (RA) cases and 335 controls. Right panel depicts the difference between cases and controls. **B)** Left panel: Barplots of the average weight proportions of each major blood cell subtype according to FACS and EpiDISH in 6 whole blood samples from Koestler et al. Root Mean Square Error (RMSE) is given. Right panel: Scatterplot of the corresponding estimated cell fractions vs the FACS estimates for all 6 samples and each cell subtype. R^2^ values (rounded to two significant digits) for each cell subtype are given, as well as the average value over all cell subtypes. **C)** Scatterplots comparing the estimated cell fractions using EpiDISH to the true known fractions for 12 experimentally reconstructed “whole blood” samples, where the mixing proportions were known. One scatterplot is shown for each blood cell subtype used in the reconstructions. In each case, we give the RMSE and R^2^ value (latter values have been rounded to two significant digits).

### EpiDISH correctly infers blood cell subtype proportions in reconstructed whole blood samples

Flow cytometric estimates of blood cell subtype proportions are also subject to error. Hence, we further assessed EpiDISH in its predictions to correctly infer cell subtype proportions from 12 reconstructed whole blood samples where the exact mixing proportions are known [20] For all those cell subtypes whose proportions exhibited a reasonable dynamic range in the experimental mixtures, EpiDISH obtained R^2^ values above 0.95, with an average RMSE of 2.6%, confirming that EpiDISH can accurately quantify blood cell subtype proportions (**Fig.2C**).

### Using DHS information marginally improves the quality of the reference database

In order to demonstrate that statistical inference is improved by using relevant cell-type DHSs when constructing the reference DNAm database [21], we conducted a comparative analysis using two different references: one using DMCs that map to cell-type specific DHSs, and another where we only use DMCs (**Methods**). We performed the analysis for three different scenarios. In the first, we generated 100 random in-silico mixtures of 5 purified blood cell subtypes from Zilbauer et al [27], and compared EpiDISH’s R^2^ value for the estimated cell-type proportions between the two different reference databases. Performing this analysis for 25 different Monte Carlo runs, revealed significantly higher R^2^ values for the database that used DHS information (**Fig.3A**). We next repeated this analysis for another scenario where we generated mixtures from 3 epithelial cell-types (breast, human renal cortical and pancreas) for which DNAm profiles were available from at least 2 independent studies, and for which DHS information for each individual cell-type was also available from either the NIH Roadmap or ENCODE (**Methods**). DNAm data from 2 independent studies is necessary to separate out the process of reference construction (training) and evaluation (validation). Confirming the previous analysis, improved R^2^ values was observed for the DHS-based reference database (**Fig.3B**). Finally, we considered a third scenario where we mixed together epithelial and non-epithelial cell-types (fetal lung fibroblast-IMR90, hepatocytes and B-cells). For each of these cell-types, DHS data and DNAm profiles generated by two independent studies were available (again, in order to avoid overfitting). Confirming the previous results, R^2^ values were distinctively improved upon using DHS information (**Fig.3C**). Using RMSE as performance measure, the DHS-based reference was best in 2/3 studies (**fig.S3 in Additional File 1**). However, we note that in all cases improvements were only marginal.

**Figure-3.**
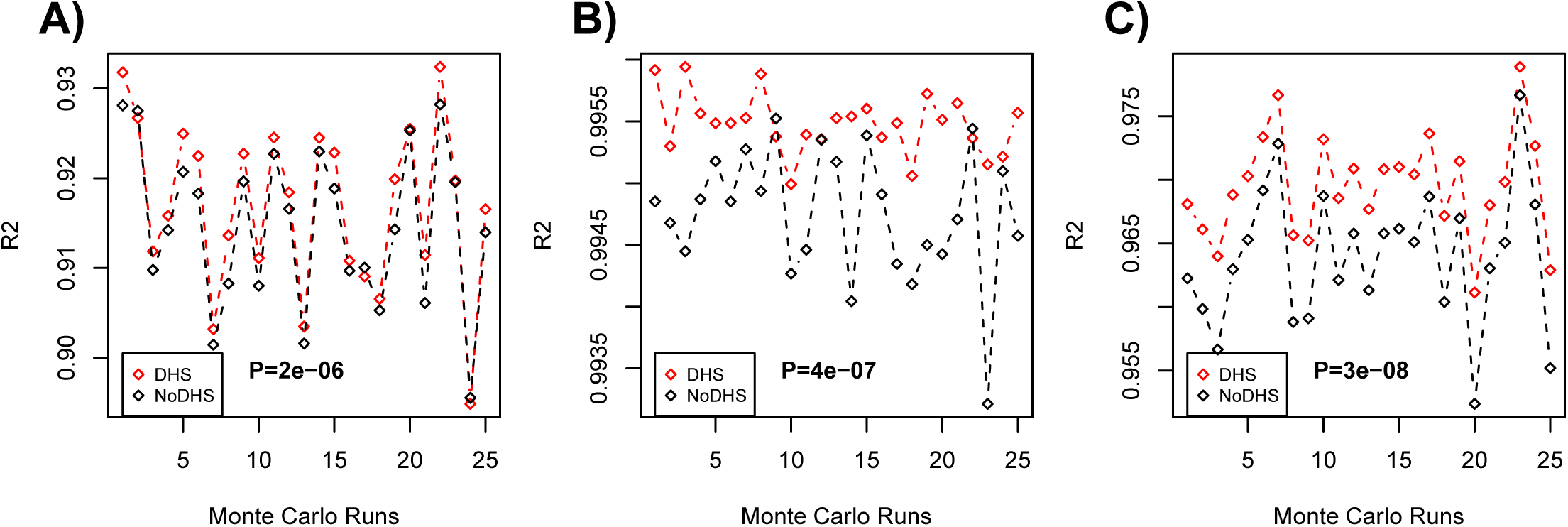
DHS information improves quality of the reference database. **A)** Using 100 in-silico mixtures of 5 purified blood cell subtypes from Zilbauer et al, we compare the R2values (R2, y-axis) of the estimated cell proportions as obtained using EpiDISH between the two different reference databases: one which uses cell-type specific DHSs to select DMCs when constructing the DNAm reference centroids (DHS), and another which only uses DMCs regardless of DHS status (noDHS). The average R2 value over all cell-types is being shown. A total of 25 different Monte Carlo runs were performed to obtain 25 average R2 values (x-axis) for each reference database. P-value is from a one-tailed paired Wilcoxon-rank sum test. **B)** As A) but now 100 in-silico generated mixtures of 3 epithelial cell subtypes (breast, renal cortical and pancreas) for which DHS data was available. **C)** As B), but now for 3 other cell subtypes (fetal lung fibroblast-IMR90, hepatocytes and B-cells) with available DHS information and DNAm profiles from two independent studies.

### EpiDISH compares favorably to other reference-based methods

Having validated EpiDISH, we next performed a detailed comparison to competing reference-based methods. Specifically, we compared EpiDISH to the linear constrained projection technique (CP) used by Houseman and others [5, 18, 19], to an algorithm called CIBERSORT (CBS), which uses Support Vector Regression and which has been shown to perform best on gene expression data [10], and finally to a non-robust variant of EpiDISH based on simple multivariate regression (LR). In order to objectively compare the 4 algorithms, we first considered the case of blood tissue, for which several datasets profiling purified blood cell subtypes were available. For each algorithm we used the same blood reference database of 333 DHS-DMCs and 7 blood cell subtypes, as constructed previously using the purified blood cell data from Reinius et al [24]. Using the known reconstructed whole blood mixtures (12 samples) from Koestler et al [20], we compared the algorithms in terms of the RMSE and R^2^ values of the estimated mixing proportions. Interestingly, we observed that robust partial correlations (EpiDISH) outperformed both CP and CIBERSORT in terms of the RMSE, with similar performance as assessed using R^2^ (**Fig.4A**). In an independent dataset of 5 purified blood cell subtypes from Zilbauer et al [27], where we generated in-silico mixtures, CP underperformed while EpiDISH/CIBERSORT performed optimally (**Fig.4B**). In order to further compare performance in the context of other cell-types, we applied all 4 algorithms to the 3 epithelial cell type and 3 epithelial/non-epithelial cell type mixture scenarios considered earlier (**Methods**). Once again, EpiDISH compared very favorably relative to the other methods, specially relative to CP which overall showed the weakest performance (**Fig.4C**, **fig.S4 in Additional File 1**). We note however that EpiDISH did not outperform CIBERSORT in one of these two studies (**fig.S4 in Additional File 1**).

**Figure-4.**
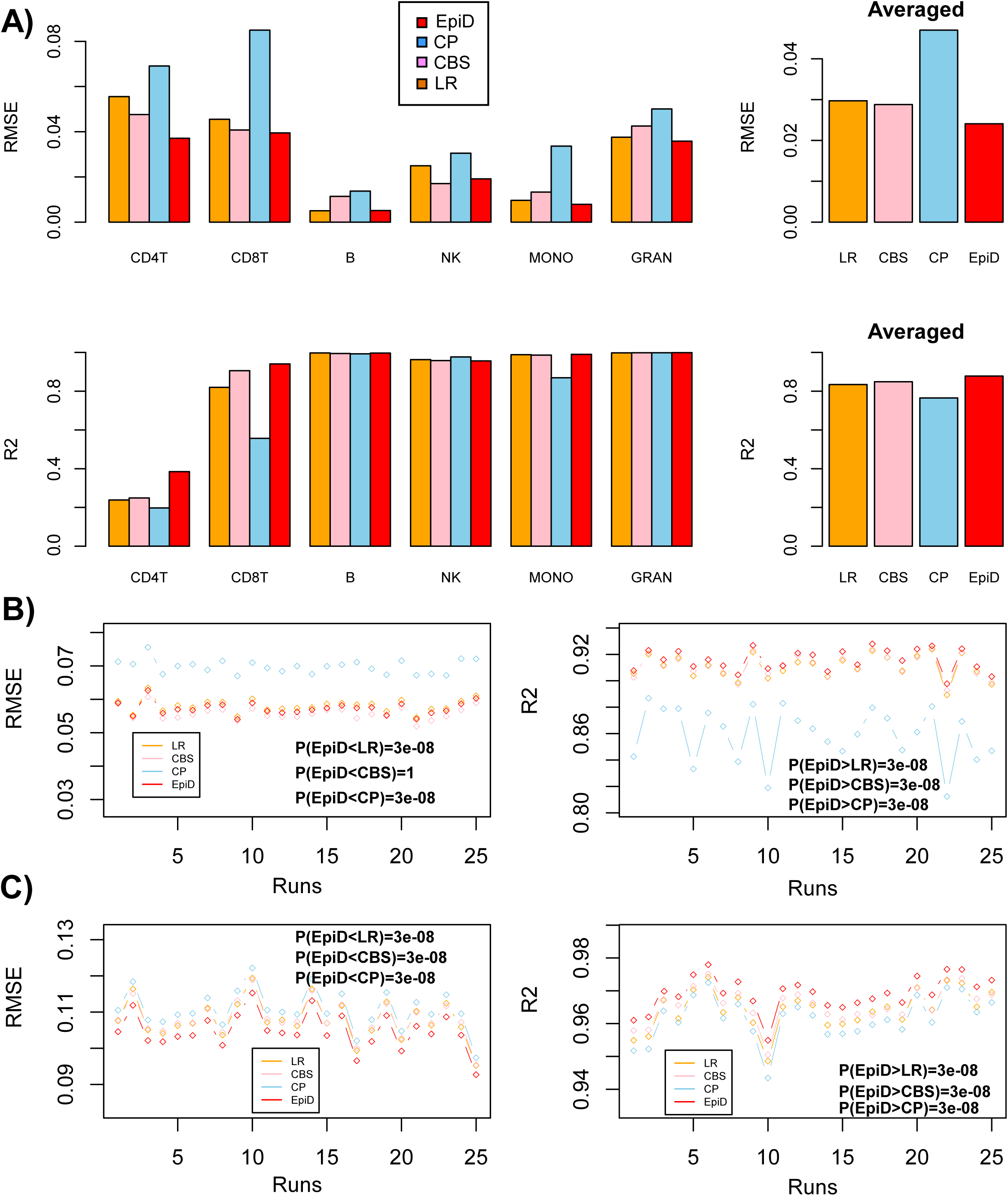
Comparison of EpiDISH to other reference-based algorithms. **A)** Barplots ofRMSE and R^2^ values for the estimated cellular proportions for the four different algorithms as applied to the reconstructed blood mixtures of Koestler et al. All estimates were obtained over the 12 reconstructed samples. Right panels show the average values over all blood cell subtypes. **B)** As A), but now for the in-silico generated mixtures of 5 blood cell subtypes from Zilbauer et al. All RMSE and R^2^ estimates were obtained over 100 randomly generated in-silico mixtures, averaged over the 5 cell-types, and this experiment was repeated a total of 25 times (“Runs”). P-values comparing EpiDISH to each of the other 3 methods are from a paired one-tailed Wilcoxon rank sum test. **C)** As B), but now for in-silico generated mixtures of 3 epithelial/non-epithelial cell types: fetal lung fibroblast, liver and B-cells. All RMSE and R^2^ estimates were obtained over 100 randomly generated in-silico mixtures, averaged over the 3 cell-types, and this experiment was repeated a total of 25 times (“Runs”). P-values comparing EpiDISH to each of the other 3 methods are from a paired one-tailed Wilcoxon rank sum test.

### Constrained Projection reveals added robustness under higher levels of noise

Since the reconstructed and in-silico mixtures from the previous analyses were generated from cell-type specific DNAm profiles that are independent from the cellular DNAm profiles used to build the reference databases, these analyses already implicitly assess the robustness of the algorithms to natural levels of variation, as encountered for instance between different labs or different experimental protocols. However, in order to improve our understanding of the noise performance characteristics of the different methods, we next investigated their relative performance under increasingly higher levels of noise (**Methods**, **Fig.5**). Adding increasing levels of noise to the reconstructed mixtures of Koestler et al, we observed that while EpiDISH was optimal for low levels, that the relative performance of other algorithms, notably constrained projection (CP), improved as noise levels increased (**Fig.5A**). We observed a similar pattern using in-silico mixtures of purified blood cell subtypes from Zilbauer et al, with EpiDISH optimal at low levels of noise, but CP emerging as the more optimal method at higher levels (**Fig.5B**). In the context of in-silico mixtures of 3 epithelial cell subtypes, we once again observed a cross-over in RMSE performance between CP and EpiDISH, with EpiDISH performing better at lower levels of noise, but CP and CIBERSORT emerging as the more optimal methods under larger levels (**Fig.5C**). In terms of R^2^, CP emerged as the best performing method in this set. CP also emerged as the better performing method (in terms of R^2^) for larger levels of noise in the context of in-silico mixtures of epithelial/non-epithelial cell subtypes (**Fig.5D**). In summary, these data indicate that the relative performance of constrained (CP) vs non-constrained (EpiDISH, CIBERSORT) approaches for estimating cell proportions in heterogeneous mixtures is dependent on cell-type and the levels of noise in the data.

**Figure-5.**
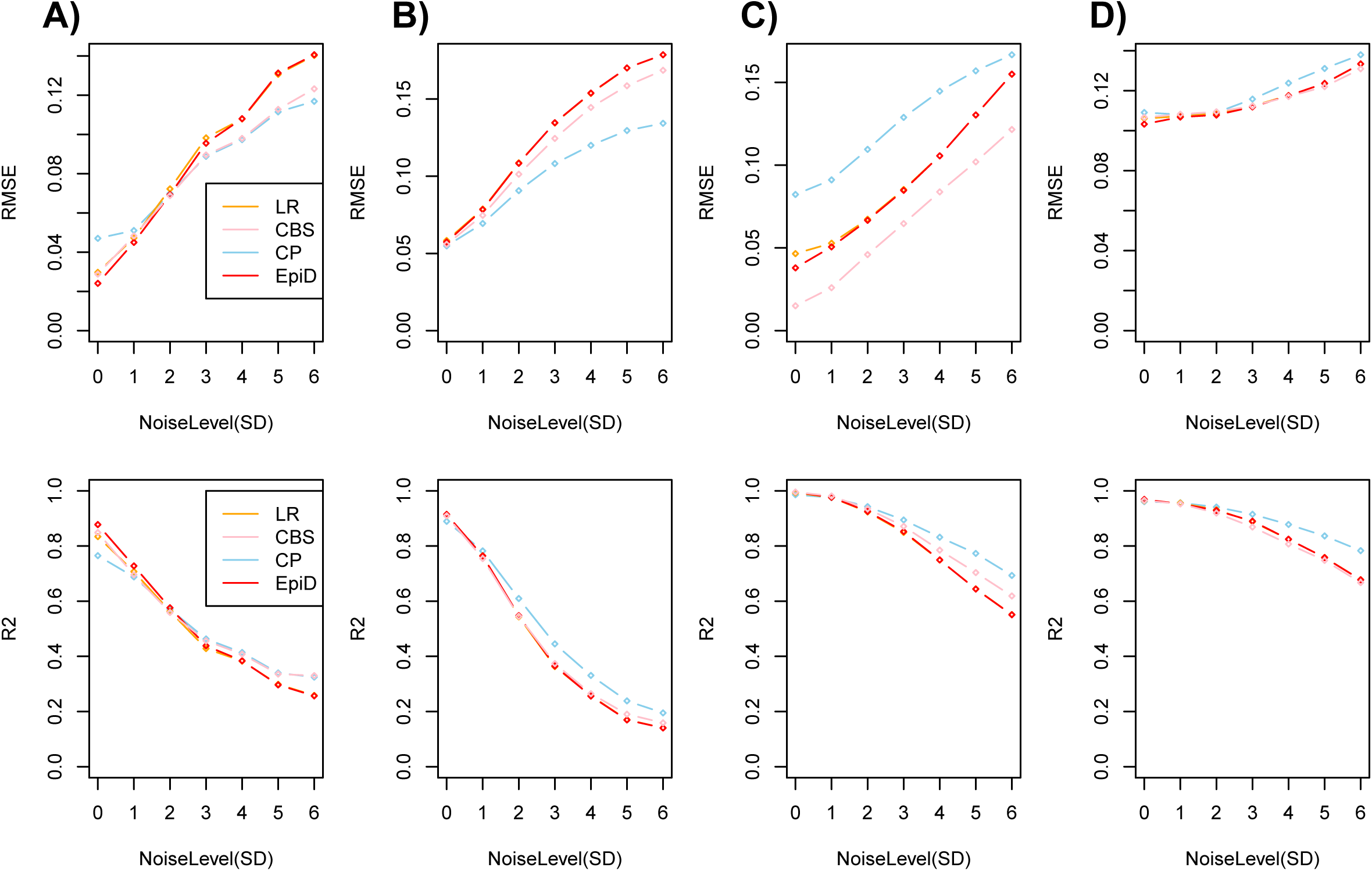
Evaluation of EpiDISH under increasing levels of noise. **A)** RMSE (upper panel) and R^2^ (lower panel) values for the estimated cellular proportions for the four different algorithms as applied to the 12 reconstructed whole blood mixtures of Koestler et al under increasing levels of noise (SD, x-axis). **B)** As A), but for 100 in-silico randomly generated mixtures of purified blood cell subtypes, profiled in Zilbauer et al. **C)** As A), but for 100 in-silico randomly generated mixtures of 3 epithelial subtypes (human mammary epithelia, human renal cortical epithelia and human pancreas). **D)** As A), but for 100 in-silico randomly generated mixtures of 3 epithelial/non-epithelial cell subtypes (fetal lung fibroblasts, liver and B-cells). In all panels A-D), RMSE and R^2^ values represent the averages over 25 repeated Monte Carlo runs.

### EpiDISH improves sensitivity in an EWAS of smoking in whole blood

In order to further validate EpiDISH and to illustrate its application to EWAS, we applied it to an EWAS of smoking in a cohort of 152 women, all aged 53, for which whole blood samples were taken and for which DNAm profiles using the Illumina 450k technology were generated [31]. Smoking in whole blood tissue provides the ideal scenario in which to test EpiDISH, since a gold-standard list of 62 smoking-associated DMCs (sDMCs), as derived from an extensive survey of independent smoking EWASs, exists [30] (**Methods, table S2 in Additional File 1**). Hence, we applied EpiDISH to our 152 samples to obtain sample-specific weights for the different blood cell subtypes. The resulting estimates were in line with what is expected for whole blood samples, with granulocytes/neutrophils making up ~50% of the samples and with lymphocytes/monocytes making up the rest (**Fig.6A**). SVD analysis confirmed that the top component of variation correlated strongly with changes in blood cell-type composition (**Fig.6B**). Not adjusting for variation in blood cell subtype composition, we observed 34 sDMCs at genome-wide significance (FDR<0.05**, table S3 in Additional File 1**, **Fig.6C**). Adjusting for cell-type proportions, as estimated using EpiDISH, we observed a doubling of sDMCs (70 sDMCs at FDR<0.05, **table S4 in Additional File 1**, Importantly, among the additional sDMCs identified with EpiDISH, there were probes that mapped to 5 genes within our gold-standard set of 15 smoking-associated genes [30, 37–41] (**Methods**, **Fig.6E**). For instance, an additional probe mapping to *AHRR* was observed after adjustment, and probes mapping to *PTK2* and *LRP5* were obtained only after adjustment (**Fig.6E**). In contrast, only one probe in the unadjusted analysis was not found after adjustment (**table S3 in Additional File 1**). To investigate this further we estimated the sensitivity for an unadjusted analysis, EpiDISH (with and without DHS-DMCs in the reference), CIBERSORT and CP, at two different FDR thresholds (FDR<0.05 and FDR<0.3) (**Table 2**). This confirmed that EpiDISH improves the sensitivity over an unadjusted analysis, although CP performed marginally better (**fig. S5 in Additional File 1**). To check that the improved sensitivity is not at the expense of a much lower specificity, we also defined a set of true negative CpGs, i.e. CpGs not associated with smoking (P>0.25) as assessed in 3 independent EWAS cohorts (**Methods**). This resulted in a true negative set of 89290 CpGs, allowing the relative specificity of the methods to be assessed. Using the same FDR < 0.05 threshold, we observed that all methods achieved comparably high relative specificity values (Specificity≈1), although CP exhibited a 3-fold higher type-1 error rate (false positive rate) than EpiDISH or CIBERSORT (**Table 2**). At a more relaxed threshold (FDR<0.3), CP exhibited a 10 times higher type-1 error rate than EpiDISH or CIBERSORT (**Table 2**). Thus, overall, EpiDISH improves inference over an unadjusted analysis and compares favorably to CP.

**Figure-6.**
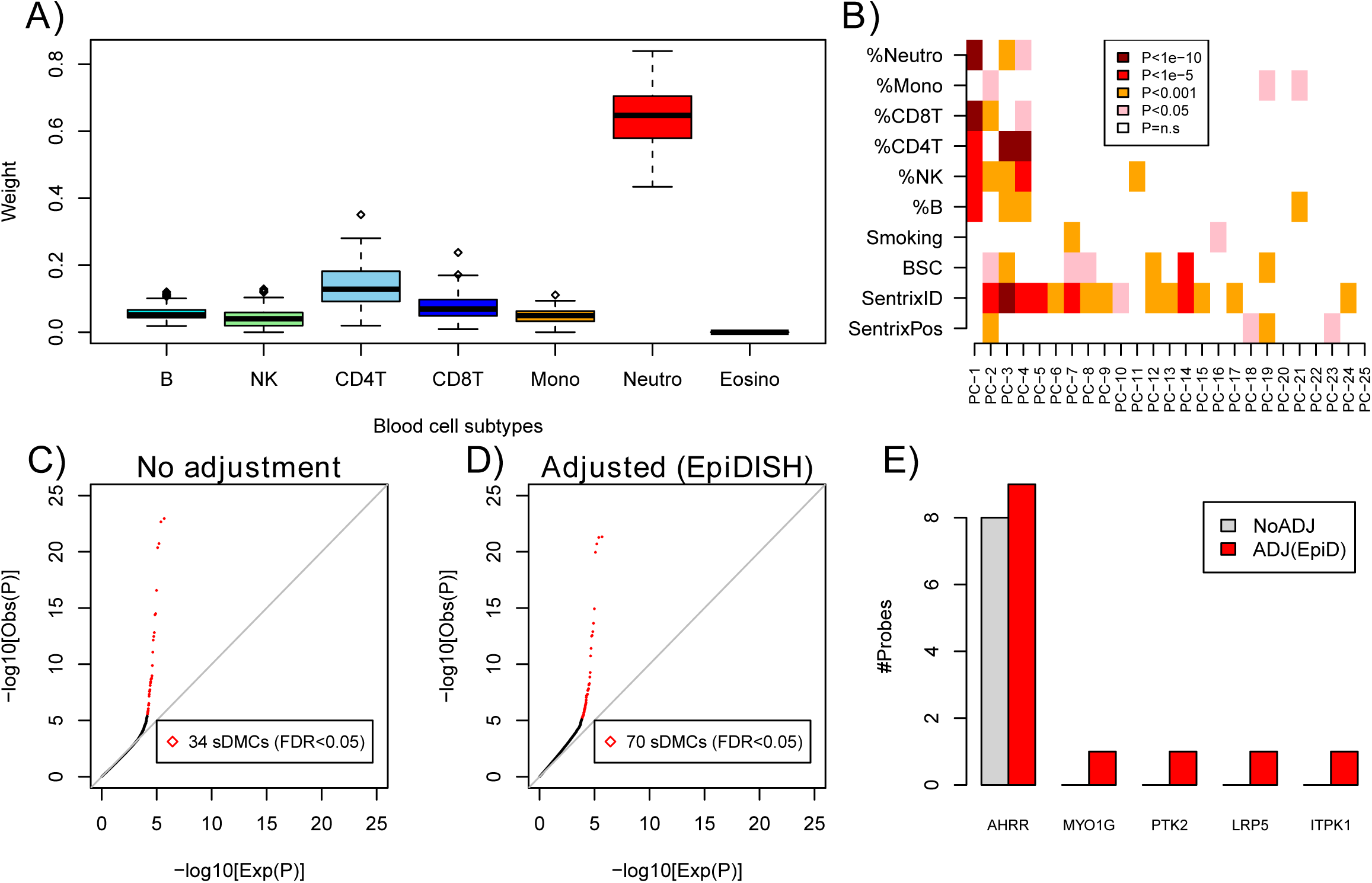
EpiDISH improves sensitivity of a smoking EWAS in blood. **A)** Distribution ofcellular proportions (weight) for the main blood cell subtypes in the 152 whole blood samples from Teschendorff et al, as inferred using EpiDISH. **B)** Heatmap of P-value associations between significant principal components (PC) and various factors, including Smoking Status, Bisulfite Conversion Efficiency (BSC), sentrix ID and position, and cellular proportion for 6 blood cell subtypes (eosinophils were not considered due to weights being effectively all zero). **C)** Quantile-quantile plot of all 450k probes passing quality control from a supervised analysis against smoking-pack-years only adjusted for sentrix ID (“No adjustment”). The number of CpGs passing an FDR<0.05 threshold are given, and defined to be smoking-associated differentially methylated CpGs (sDMCs). **D)** Quantile-quantile plot of all 450k probes passing quality control from a supervised analysis against smoking-pack-years adjusted for sentrix ID and blood cell subtype proportions as estimated using EpiDISH (“Adjusted (EpiDISH)”). The number of CpGs passing an FDR<0.05 threshold are given, and defined to be smoking-associated differentially methylated CpGs (sDMCs). **E)** Among the 34 and 70 sDMCs identified in C) and D), respectively, we indicate the numbers of these that map to a selected set of 5 well-known and validated smoking-associated genes. This subset derives from a set of 15 gold-standard smoking-associated genes, as curated by a review of the literature by Gao et al [30].

**Table 2:** Comparison of the relative sensitivity and specificity. Smoking-associated DMCs (sDMCs) were defined at FDR<0.05 and FDR<0.3 and for each of 5 methods in the 152 whole blood Illumina 450k samples from the MRC1946 birth cohort data. Methods are: NoADJ=no-adjustment, EpiD(noDHS)=adjustment with RPC using reference with no DHS info, EpiD=adjustment with RPC and a reference with DHS info, CBS=adjustment with CIBERSORT, CP=adjustment with constrained projection. Sensitivity was estimated as the fraction of the 62 gold-standard true positives (TP) found among the sDMCs. In the case of specificity, this was estimated as the fraction of true negatives (TN) among sDMCs measured relative to a gold-standard set of 89290 true negative (TN) CpGs.

| **nTP=62, nTN=89290** | **NoADJ** | **EpiD(noDHS)** | **EpiD** | **CBS** | **CP** |
| --- | --- | --- | --- | --- | --- |
| **FDR<0.05** |  |  |  |  |  |
| **sDMCs** | 34 | 67 | 70 | 58 | 152 |
| **\| TP ∩ sDMC \|** | 31 | 34 | 35 | 36 | 37 |
| **Sensitivity** | 0.56 | 0.55 | 0.56 | 0.58 | 0.60 |
| **FP=\| TN ∩ sDMC \|** | 0 | 3 | 3 | 0 | 11 |
| **Specificity** | 1 | 0.999966 | 0.999966 | 1 | 0.999877 |
| **Empirical FDR** | <0.03 | 0.045 | 0.043 | <0.02 | 0.072 |
| **FDR<0.3** |  |  |  |  |  |
| **sDMCs** | 67 | 544 | 802 | 259 | 10457 |
| **\| TP ∩ sDMC \|** | 40 | 50 | 52 | 44 | 54 |
| **Sensitivity** | 0.64 | 0.81 | 0.84 | 0.71 | 0.87 |
| **FP=\| TN ∩ sDMC \|** | 1 | 73 | 121 | 31 | 1735 |
| **Specificity** | 0.99998 | 0.9992 | 0.998 | 0.9996 | 0.98 |
| **Empirical FDR** | 0.01 | 0.13 | 0.15 | 0.12 | 0.17 |

## Discussion

The aim of this study was to compare different reference-based methods in order to establish whether there is an optimal approach. This is particularly pertinent given that only one reference-based approach (Houseman’s algorithm) has been applied to DNA methylation data. As our study has demonstrated, Houseman’s algorithm, which relies on the CP technique, is in fact not the most robust method. While we did not identify a single method which always outperformed all others, we found that non-constrained techniques such as partial correlations and support vector regression provided a more robust inference framework than CP, particularly for more realistic noise levels. This is consistent with the results obtained by Newman et al on gene expression data [10]. In our study, CP only emerged as the optimal choice if data was subjected to a large amount of noise, typically larger than what is encountered in real data. Hence, our study advances upon the state-of-the-art, and proposes the use of RPC or CIBERSORT as a more robust means for inferring cell-fractions in complex tissues.

Another important novel insight of our study is that inference can be improved, albeit only marginally, by incorporating cell-type specific DHS information when constructing the reference DNAm database. We demonstrated this not only in the context of blood tissue, but also for mixtures involving epithelial cell subtypes. To understand why the improvement is only marginal, we note that supervised selection of DMCs from a training set, for instance, by selecting DMCs that show big (i.e. 80% or over) differences in DNAm between cell-types, will almost always identify true DMCs. Supervised selection tends to favour DMCs with larger differences in average DNAm which serve better for the statistical deconvolution problem. Thus, while prior biological knowledge can help remove a few false positives, this leads to only a relatively minor improvement in the quality of the inference. It will be interesting to explore if further improvements are possible, for instance, using predicted cell-type specific DMCs [42], although we anticipate that such prior information will turn out to be more fruitful in the context of “semi-reference-free” approaches such as RUV [43].

We further showcased EpiDISH in an EWAS of smoking, where it was found to identify twice as many smoking-associated DMCs than an unadjusted analysis. Importantly, some of the additional sDMCs mapped to genes (e.g. *AHRR, PTK2, MYO1G, LRP5*) which are well-known to be associated with differential methylation in smokers [30], thus confirming that EpiDISH leads to an increase in sensitivity. Although similar improvements were possible with CIBERSORT and CP, CP suffered from a higher FDR. Thus, our detailed analysis on experimental and in-silico generated mixtures, which suggests improved modelling with a non-constrained technique such as RPCs, appears to also hold true on real data, although we caution that analyses in more smoking-EWAS will be needed to reach a robust conclusion.

Our analysis focused entirely on Illumina 450k data. While this may be seen as a limitation, in light of the recent arrival of the newer 850k version [44], the overwhelming majority of the 450k probes are present in the new beadarray version, rendering the extension to 850k data relatively trivial. Likewise, given the relatively good agreement between Illumina 450k and WGBS/RRBS data [45], and given that in both cases DNAm data can be treated in the beta-value basis, we would envisage that application of EpiDISH will carry over to such data, although studies demonstrating this will be needed.

## Conclusions

In summary, we have presented a novel reference-based algorithm, EpiDISH, for in-silico deconvolution of DNA methylation data, which compares very favorably in relation to the current gold-standard. We recommend the use of EpiDISH for dissection of intra-sample heterogeneity in EWAS, and to this purpose we provide the community with an R-package (EpiDISH), freely available from *https://github.com/sjczheng/EpiDISH.*

## Abbreviations

DMC: differentially methylated CpG
EWAS: Epigenome-Wide-Association Study
DNAm: DNA methylation
FDR: false discovery rate
RUV: Removing Unwanted Variation
DHS: DNAse Hypersensitive Site
sDMC: smoking-associated differentially methylated CpG

## Declarations

### Ethics approval and consent to participate

Not applicable.

### Consent to publish

Not Applicable

## Acknowledgements

None

## Availability of data and materials

All data analysed in this study is publicly available from repositories as indicated in Methods. EpiDISH is freely available as an R-package from *github: https://github.com/sjczheng/EpiDISH*.

## Authors’ contributions

AET performed most of the statistical analyses and wrote the manuscript. SCZ contributed to data analysis and software-testing. CEB and SB provided data. Study was conceived by AET with input from CEB and SB.

## Description of Additional Files

**Additional File 1:** Contains all Supplementary Figures and Supplementary Tables plus their captions:

**fig. S1: Clustering validation of reference blood database.**

**fig. S2: Validation of EpiDISH.**

**fig. S3: Improvement of inference using DHS data in 3 independent data sets.**

**fig. S4: Comparison of reference-based methods.**

**fig. S5: Comparative performance of reference-based methods in an EWAS of smoking.**

**table S1: The blood reference database.**

**table S2: Gold-standard list of 62 smoking-associated DMCs (sDMCs).**

**table S3: Smoking-associated DMCs with no adjustment for cell-type composition.**

**table S4: Smoking-associated DMCs as obtained using EpiDISH.**

**Additional File 2:** An R-script implementing EpiDISH with either robust partial correlations (RPC), CIBERSORT (CBS) or Constrained Projection (CP).

## References

[1] Alizadeh AA, Aranda V, Bardelli A, Blanpain C, Bock C, Borowski C, Caldas C, Califano A, Doherty M, Elsner M, et al: Toward understanding and exploiting tumor heterogeneity. Nat Med 2015, 21:846–853.

[2] Liu Y, Aryee MJ, Padyukov L, Fallin MD, Hesselberg E, Runarsson A, Reinius L Acevedo N, Taub M, Ronninger M, et al: Epigenome-wide association data implicate DNA methylation as an intermediary of genetic risk in rheumatoid arthritis. Nat Biotechnol 2013, 31:142–147.

[3] Jaffe AE Irizarry RA: Accounting for cellular heterogeneity is critical in epigenome-wide association studies. Genome Biol 2014, 15:R31.

[4] Rakyan VK, Down TA, Balding DJ, Beck S: Epigenome-wide association studies for common human diseases. Nat Rev Genet 2011, 12:529–541.

[5] Houseman EA, Accomando WP, Koestler DC, Christensen BC, Marsit CJ, Nelson HH, Wiencke JK, Kelsey KT: DNA methylation arrays as surrogate measures of cell mixture distribution. BMC Bioinformatics 2012, 13:86.

[6] Zou J, Lippert C, Heckerman D, Aryee M, Listgarten J: Epigenome-wide association studies without the need for cell-type composition. Nat Methods 2014.

[7] Houseman EA, Kelsey KT, Wiencke JK, Marsit CJ: Cell-composition effects in the analysis of DNA methylation array data: a mathematical perspective. BMC Bioinformatics 2015, 16:95.

[8] Houseman EA, Molitor J, Marsit CJ: Reference-free cell mixture adjustments in analysis of DNA methylation data. Bioinformatics 2014, 30:1431–1439.

[9] Zheng X, Zhao Q, Wu HJ, Li W, Wang H, Meyer CA, Qin QA, Xu H, Zang C, Jiang P, et al: MethylPurify: tumor purity deconvolution and differential methylation detection from single tumor DNA methylomes. Genome Biol 2014, 15:419.

[10] Newman AM, Liu CL, Green MR, Gentles AJ, Feng W, Xu Y, Hoang CD, Diehn M, Alizadeh AA: Robust enumeration of cell subsets from tissue expression profiles. Nat Methods 2015, 12:453–457.

[11] Yoshihara K, Shahmoradgoli M, Martinez E, Vegesna R, Kim H, Torres-Garcia W, Trevino V, Shen H, Laird PW, Levine DA, et al: Inferring tumour purity and stromal and immune cell admixture from expression data. Nat Commun 2013, 4:2612.

[12] Teschendorff AE, Menon U, Gentry-Maharaj A, Ramus SJ, Gayther SA, Apostolidou S, Jones A, Lechner M, Beck S, Jacobs IJ, Widschwendter M: An epigenetic signature in peripheral blood predicts active ovarian cancer. PLoS One 2009, 4:e8274.

[13] Langevin SM, Koestler DC, Christensen BC, Butler RA, Wiencke JK, Nelson HH, Houseman EA, Marsit CJ, Kelsey KT: Peripheral blood DNA methylation profiles are indicative of head and neck squamous cell carcinoma: an epigenome-wide association study. Epigenetics 2012, 7:291–299.

[14] Vogt H, Hofmann B, Getz L: The new holism: P4 systems medicine and the medicalization of health and life itself. Med Health Care Philos 2016.

[15] Timp W, Bravo HC, McDonald OG, Goggins M, Umbricht C, Zeiger M, Feinberg AP, Irizarry RA: Large hypomethylated blocks as a universal defining epigenetic alteration in human solid tumors. Genome Medicine 2014, 6.

[16] Teschendorff AE, Gao Y, Jones A, Ruebner M, Beckmann MW, Wachter DL, Fasching PA, Widschwendter M: DNA methylation outliers in normal breast tissue identify field defects that are enriched in cancer. Nat Commun 2016, 7:10478.

[17] Roadmap Epigenomics C, Kundaje A, Meuleman W, Ernst J, Bilenky M, Yen A, Kheradpour P, Zhang Z, Wang J, et al: Integrative analysis of 111 reference human epigenomes. Nature 2015, 518:317–330.

[18] Koestler DC, Christensen B, Karagas MR, Marsit CJ, Langevin SM, Kelsey KT, Wiencke JK, Houseman EA: Blood-based profiles of DNA methylation predict the underlying distribution of cell types: a validation analysis. Epigenetics 2013, 8:816–826.

[19] Accomando WP, Wiencke JK, Houseman EA, Nelson HH, Kelsey KT: Quantitative reconstruction of leukocyte subsets using DNA methylation. Genome Biol 2014, 15:R50.

[20] Koestler DC, Jones MJ, Usset J, Christensen BC, Butler RA, Kobor MS, Wiencke JK, Kelsey KT: Improving cell mixture deconvolution by identifying optimal DNA methylation libraries (IDOL). BMC Bioinformatics 2016, 17:120.

[21] Thurman RE, Rynes E, Humbert R, Vierstra J, Maurano MT, Haugen E, Sheffield NC, Stergachis AB, Wang H, Vernot B, et al: The accessible chromatin landscape of the human genome. Nature 2012, 489:75–82.

[22] Dunham I, Kundaje A, Aldred SF, Collins PJ, Davis CA, Doyle F, Epstein CB, Frietze S, Harrow J, Kaul R, et al: An integrated encyclopedia of DNA elements in the human genome. Nature 2012, 489:57–74.

[23] Gerstein MB, Kundaje A, Hariharan M, Landt SG, Yan KK, Cheng C, Mu XJ, Khurana E, Rozowsky J, Alexander R, et al: Architecture of the human regulatory network derived from ENCODE data. Nature 2012, 489:91–100.

[24] Reinius LE, Acevedo N, Joerink M, Pershagen G, Dahlen SE, Greco D, Soderhall C, Scheynius A, Kere J: Differential DNA methylation in purified human blood cells: implications for cell lineage and studies on disease susceptibility. PLoS One 2012, 7:e41361.

[25] Smyth GK: Linear models and empirical bayes methods for assessing differential expression in microarray experiments. Stat Appl Genet Mol Biol 2004, 3:Article3.

[26] Slieker RC, Bos SD, Goeman JJ, Bovee JV, Talens RP, van der Breggen R, Suchiman HE, Lameijer EW, Putter H, van den Akker EB, et al: Identification and systematic annotation of tissue-specific differentially methylated regions using the Illumina 450k array. Epigenetics Chromatin 2013, 6:26.

[27] Zilbauer M, Rayner TF, Clark C, Coffey AJ, Joyce CJ, Palta P, Palotie A, Lyons PA, Smith KG: Genome-wide methylation analyses of primary human leukocyte subsets identifies functionally important cell-type-specific hypomethylated regions. Blood 2013, 122:e52–60.

[28] Lowe R, Overhoff MG, Ramagopalan SV, Garbe JC, Koh J, Stampfer MR, Beach DH, Rakyan VK, Bishop CL: The senescent methylome and its relationship with cancer, ageing and germline genetic variation in humans. Genome Biol 2015, 16:194.

[29] Nazor KL, Altun G, Lynch C, Tran H, Harness JV, Slavin I, Garitaonandia I, Muller FJ, Wang YC, Boscolo FS, et al: Recurrent variations in DNA methylation in human pluripotent stem cells and their differentiated derivatives. Cell Stem Cell 2012, 10:620–634.

[30] Gao X, Jia M, Zhang Y, Breitling LP, Brenner H: DNA methylation changes of whole blood cells in response to active smoking exposure in adults: a systematic review of DNA methylation studies. Clin Epigenetics 2015, 7:113.

[31] Teschendorff AE, Yang Z, Wong A, Pipinikas CP, Jiao Y, Jones A, Anjum S, Hardy R, Salvesen HB, Thirlwell C, et al: Correlation of Smoking-Associated DNA Methylation Changes in Buccal Cells With DNA Methylation Changes in Epithelial Cancer. JAMA Oncol 2015, 1:476–485.

[32] Tsaprouni LG, Yang TP, Bell J, Dick KJ, Kanoni S, Nisbet J, Vinuela A, Grundberg E, Nelson CP, Meduri E, et al: Cigarette smoking reduces DNA methylation levels at multiple genomic loci but the effect is partially reversible upon cessation. Epigenetics 2014, 9:1382–1396.

[33] Aryee MJ, Jaffe AE, Corrada-Bravo H, Ladd-Acosta C, Feinberg AP, Hansen KD, Irizarry RA: Minfi: a flexible and comprehensive Bioconductor package for the analysis of Infinium DNA methylation microarrays. Bioinformatics 2014, 30:1363–1369.

[34] Teschendorff AE, Marabita F, Lechner M, Bartlett T, Tegner J, Gomez-Cabrero D, Beck S: A beta-mixture quantile normalization method for correcting probe design bias in Illumina Infinium 450 k DNA methylation data. Bioinformatics 2013, 29:189–196.

[35] Lowe R, Rakyan VK: Marmal-aid–a database for Infinium HumanMethylation450. BMC Bioinformatics 2013, 14:359.

[36] Bernstein BE, Stamatoyannopoulos JA, Costello JF, Ren B, Milosavljevic A, Meissner A, Kellis M, Marra MA, Beaudet AL, Ecker JR, et al: The NIH Roadmap Epigenomics Mapping Consortium. Nat Biotechnol 2010, 28:1045–1048.

[37] Zeilinger S, Kuhnel B, Klopp N, Baurecht H, Kleinschmidt A, Gieger C, Weidinger S, Lattka E, Adamski J, Peters A, et al: Tobacco smoking leads to extensive genome-wide changes in DNA methylation. PLoS One 2013, 8:e63812.

[38] Shenker NS, Ueland PM, Polidoro S, van Veldhoven K, Ricceri F, Brown R, Flanagan JM, Vineis P: DNA methylation as a long-term biomarker of exposure to tobacco smoke. Epidemiology 2013, 24:712–716.

[39] Zhang Y, Yang R, Burwinkel B, Breitling LP, Brenner H: F2RL3 methylation as a biomarker of current and lifetime smoking exposures. Environ Health Perspect 2014, 122:131–137.

[40] Breitling LP, Yang R, Korn B, Burwinkel B, Brenner H: Tobacco-smoking-related differential DNA methylation: 27K discovery and replication. Am J Hum Genet 2011, 88:450–457.

[41] Besingi W, Johansson A: Smoke-related DNA methylation changes in the etiology of human disease. Hum Mol Genet 2014, 23:2290–2297.

[42] Ernst J, Kellis M: Large-scale imputation of epigenomic datasets for systematic annotation of diverse human tissues. Nat Biotechnol 2015, 33:364–376.

[43] Gagnon-Bartsch JA, Speed TP: Using control genes to correct for unwanted variation in microarray data. Biostatistics 2012, 13:539–552.

[44] Moran S, Arribas C, Esteller M: Validation of a DNA methylation microarray for 850,000 CpG sites of the human genome enriched in enhancer sequences. Epigenomics 2016, 8:389–399.

[45] Titus AJ, Houseman EA, Johnson KC, Christensen BC: methyLiftover: cross-platform DNA methylation data integration. Bioinformatics 2016.

